# Accumulation of ibuprofen in endemic amphipods of Lake Baikal

**DOI:** 10.1101/2025.09.04.673924

**Authors:** Tamara Y. Telnova, Maria M. Morgunova, Sophie S. Shashkina, Maria E. Dmitrieva, Victoria N. Shelkovnikova, Olga E. Lipatova, Ekaterina V. Malygina, Natalia A. Imidoeva, Alexander Y. Belyshenko, Tatyana N. Vavilina, Arkadii N. Matveev, Evgenia A. Misharina, Denis V. Axenov-Gribanov

**Affiliations:** Bioorganics Research and Educational Center, Irkutsk State University, Irkutsk, Russia; UNESCO Chair on Water Resources, Irkutsk State University, Irkutsk, Russia; Institute of biological sciences, Irkutsk State University, Irkutsk, Russia

**Keywords:** amphipods, pharmaceutical pollution, ibuprofen, Lake Baikal, accumulation, endemics

## Abstract

Pharmaceutical pollutants, including ibuprofen, are now ubiquitously detected in global aquatic ecosystems, exerting significant negative ecological impacts. Lake Baikal organisms have been documented to accumulate ibuprofen and related contaminants. This study quantified ibuprofen concentrations within Lake Baikal’s endemic amphipod fauna using high-performance liquid chromatography coupled with mass spectrometry (HPLC-MS). We analyzed specimens representing key ecological groups across the genera *Eulimnogammarus, Brandtia, Ommatogammarus*, and *Pallasea*. Ibuprofen concentrations ranged from 4.19 ng/g to 1151.32 ng/g (wet weight), confirming consistent contamination in both littoral and deep-water endemic amphipod populations.

Crucially, our data provide the first evidence suggesting amphipods, or their associated symbiotic microbiota, may metabolize ibuprofen. Interspecific accumulation patterns were identified, with *Eulimnogammarus* sp. and *Brandtia* sp. exhibiting distinct profiles. Furthermore, accumulation was significantly higher during spring compared to autumn samples. A negative correlation emerged between ibuprofen concentration and amphipod body mass within species. Several populations contained non-detectable levels.

These findings demonstrate that endemic amphipods within Lake Baikal’s natural environment are exposed to and bioaccumulate the pharmaceutical pollutant ibuprofen, exhibiting species-specific, seasonal, and allometric variation in uptake.

## INTRODUCTION

Pharmaceuticals are indispensable in modern healthcare and daily life, safeguarding public health while enhancing quality and longevity. Non-steroidal anti-inflammatory drugs (NSAIDs) constitute a major therapeutic class, extensively employed in human and veterinary medicine for their analgesic, antipyretic, and anti-inflammatory properties (Parolini, 2020).

Global accessibility and prevalence drive escalating NSAID demand and production. However, regulatory oversight remains limited, with most NSAIDs available over-the-counter (Jan-Roblero & Cruz-Maya, 2023). An estimated 30 million individuals use NSAIDs daily, exceeding 300 million users annually worldwide. This high consumption poses significant environmental challenges (Nieto et al., 2017; Jurado et al., 2021) due to incomplete drug metabolism in organisms, inadequate disposal practices and insufficient removal by wastewater treatment facilities. Following ingestion, a substantial fraction of NSAIDs is excreted unmetabolized or as bioactive compounds (Marchlewicz et al., 2015), ultimately entering aquatic ecosystems. The resultant influx of pharmaceutical pollutants exerts detrimental effects on freshwater biota. Monitoring studies consistently detect NSAIDs— including acetylsalicylic acid, paracetamol (acetaminophen), diclofenac, ketoprofen, and naproxen— in freshwater systems globally (Tyumina et al., 2020), underscoring their pervasive environmental presence.

Notably, ibuprofen stands out among NSAIDs due to its extensive clinical use and inclusion on the World Health Organization’s List of Essential Medicines (Chopra & Kumar, 2020). However, its high global consumption—with annual production exceeding 300,000 tonnes (Marchlewicz et al., 2015)—presents a significant environmental risk. Ibuprofen exhibits limited aqueous solubility and undergoes incomplete metabolism in humans. Following excretion, it enters the environment either unmetabolized or as biotransformation products, primarily hydroxyibuprofen, carboxyibuprofen, carboxyhydratropic acid (Chopra & Kumar, 2020), and 4-isobutylcatechol (Larsen, 2019). These metabolites often undergo further hydrolysis and are frequently more toxic to aquatic organisms than the parent compound (Chopra & Kumar, 2020).

Global monitoring studies confirm ibuprofen’s pervasive presence in aquatic systems. Reported concentrations in surface waters range from 0.98 to 1.417 µg/L across diverse regions including Canada, France, China, Greece, Korea, Taiwan, and Uganda (Kim et al., 2009; Vulliet, 2011; Almeida et al., 2013; Luo et al., 2014; Nantaba et al., 2020). Conversely, groundwater studies in Europe report lower but still detectable levels, ranging from 3 to 395 ng/L (Luo et al., 2014), underscoring its environmental persistence.

Furthermore, ibuprofen and its metabolites exert demonstrable toxicity on aquatic organisms (Das et al., 2019). Field studies confirm bioaccumulation across taxa: *Hydropsyche* spp. caddisflies in Spain’s Segre River contained 184 ng/g ibuprofen (Huerta et al., 2015), while *Gammarus fossarum* amphipods downstream of a wastewater treatment plant in southern France showed concentrations of 60.6–105.4 ng/g (Berlioz-Barbier et al., 2014). In urbanized Chinese rivers, ibuprofen was detected in phytoplankton (14.5–35.8 ng/g), zooplankton (20.9–48.9 ng/g), and benthic invertebrates (freshwater shrimp, mussels, snails: 4.8–11.6 ng/g) (Yang et al., 2020). Laboratory studies indicate adverse effects occur at concentrations typically orders of magnitude higher than environmental levels (10–100 mg/L) (Pajić et al., 2023). Documented toxicity spans diverse aquatic taxa, including echinoderms (*Asterias rubens, Psammechinus miliaris*), polychaetes (*Arenicola marina*), microalgae (*Navicula* sp., *Chlorella vulgaris, Acutodesmus obliquus, Chlamydomonas reinhardtii, Nannochloropsis limnetica*), and crustaceans (*Daphnia magna*) (Grzesiuk, 2016; Geiger, 2016; Du et al., 2016; Zanuri, 2017; Ding et al., 2019).

These findings underscore that ibuprofen contamination poses a potential yet understudied threat to ancient aquatic ecosystems. Lake Baikal – a tectonic-formed UNESCO World Heritage site in southeastern Siberia – represents a critical case study as Earth’s deepest lake (Moore et al., 2009, 2019; Rusinek et al., 2012) and one of its most ancient, with recent geological evidence indicating an age exceeding 60 million years (Matz & Efimova, 2017). This evolutionary cradle harbors >2,600 animal species exhibiting exceptional endemism (Timoshkin et al., 2016), heightening vulnerability to anthropogenic pollutants like ibuprofen.

As in all aquatic ecosystems, pollutants in Lake Baikal undergo trophic transfer via nekton, plankton, and benthos (Rusinek et al., 2015). Among endemic benthic taxa, amphipods (Amphipoda, Crustacea) represent a hyperdiverse and ubiquitous group occupying all depth zones and substrate types (Takhteev, 2019). This ecological dominance positions them as sentinel organisms for early pollutant exposure.

Critically, recent research confirms Baikal amphipods bioaccumulate multiple pharmaceuticals—including acetylsalicylic acid, paracetamol, tetracycline antibiotics, and notably ibuprofen (Telnova et al., 2024). The persistent detection of ibuprofen, a compound with established ecotoxicity, signals direct risks to endemic species and broader lake ecosystem integrity. To address key knowledge gaps, this study quantifies species- and population-specific ibuprofen concentrations in endemic Baikal amphipods and evaluates seasonal accumulation dynamics.

## MATERIALS AND METHODS

The current study focused on adult amphipods spanning key ecological niches within Lake Baikal. Specimens included littoral species (*Eulimnogammarus cyaneus* (Dybowsky, 1874), *E. verrucosus* (Gerstfeldt, 1858), and unidentified *Eulimnogammarus* specimens), free-living sublittoral species (*Brandtia* sp. (Bate, 1862)), sublittoral benthic species (*Pallasea* sp. (Bate, 1862)), and deep-water species *Ommatogammarus flavus* (Dybowsky, 1874). Detailed ecological and physiological characterization of these taxa was well described in earlier studies (Axenov-Gribanov et al., 2016; Shirokova et al., 2024; Drozdova et al., 2025).

Amphipods related to species *E. verrucosus* were collected from littoral zones at: Kultuk, Listvyanka, Bolshoye Goloustnoye, Ust-Barguzin, and Buguldeika settlements (South and Middle Baikal) during spring 2023. Additional specimens were obtained from the Angara River in Irkutsk city. The place of sampling in Angara was located upstream of municipal wastewater treatment facilities. Listvyanka populations of *E. verrucosus* were sampled seasonally (spring and autumn 2023).

Specimens of *E. cyaneus* were obtained from littoral zone of Angara River during spring 2023. Amphipods of *Brandtia* sp. were collected concurrently at two sites: the Angara River and Listvyanka settlement. *O. flavus* was collected from the profundal zone (70 m depth) near Buguldeika settlement using hydrobiological traps (spring 2023). All specimens were collected with hydrobiological nets except where noted.

All specimens were immediately transferred to 2 mL plastic Eppendorf tubes, flash-frozen in liquid nitrogen, and stored at −196°C. Sampling locations are shown in Figure 1. Lake Baikal experiences intense recreational pressure as a major tourism hub, with all sampling sites situated in highly frequented destinations (Aleksandrova et al., 2021). This anthropogenic context heightens contamination risks from pharmaceutical residues in nearshore ecosystems.

**Fig. 1.**
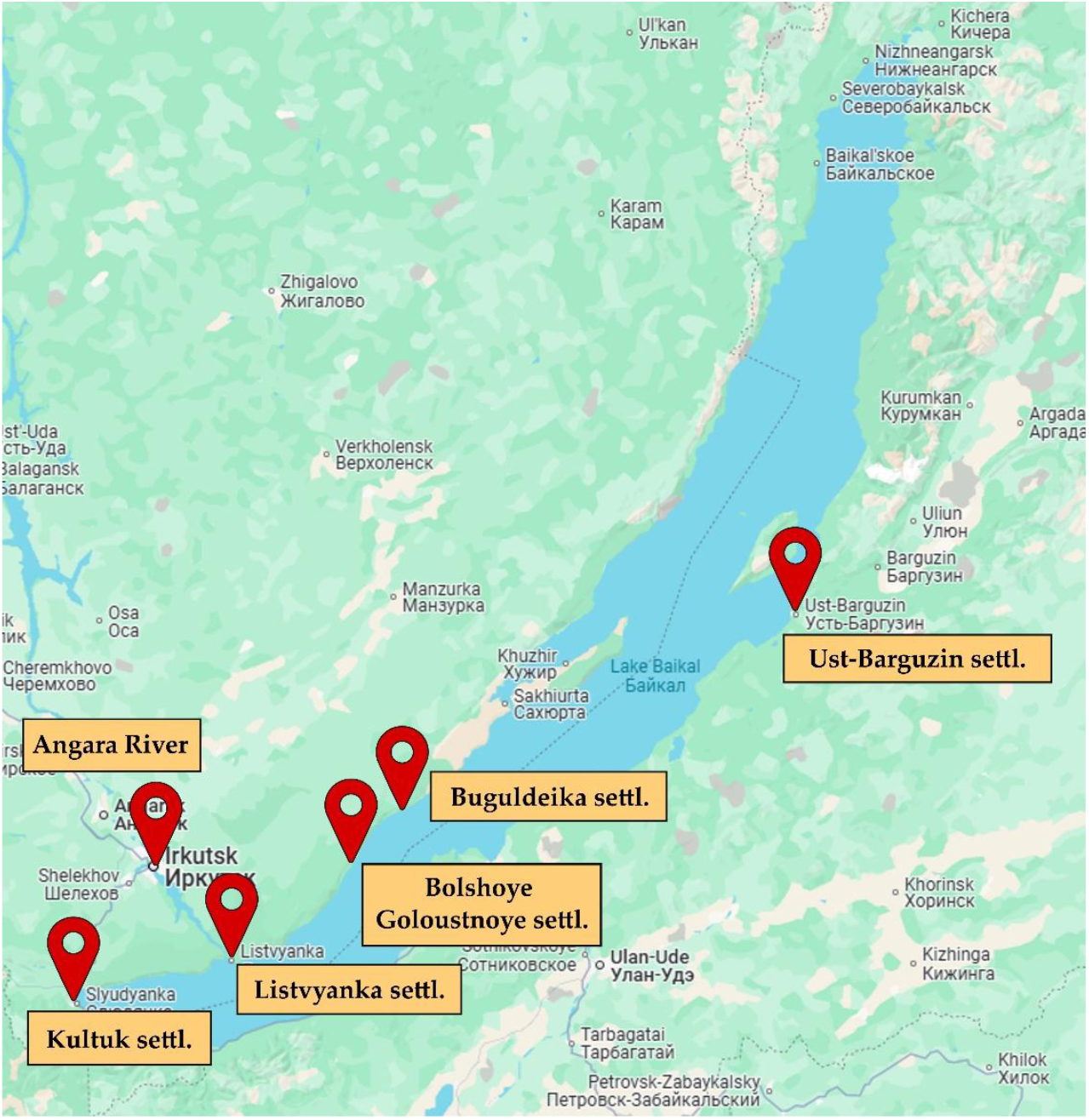
Map of amphipod sampling sites, Lake Baikal

The study included two stages, and involved qualitative and quantitative assessments of ibuprofen content in amphipods. Each amphipod was weighed individually on an analytical balance (Sartorius CE224-C, St. Petersburg, Russia) and homogenized three times in a vibratory ball mill (BABRx1, Mycotech, Irkutsk, Russia) with acetonitrile (Telnova et al., 2024). After each grinding cycle, samples were briefly centrifuged at 1,000 rpm for 1 min (Armed LC-04B, St. Petersburg, Russia) to separate supernatant and debris. Supernatant was transferred into new 2-mL Eppendorf tubes, and debris was re-extracted. After complete extraction, samples were brought to a final volume of 4 mL with acetonitrile. The tubes were centrifuged (Armed LC-04B, St. Petersburg, Russia) for 10 minutes at 3,000 rpm. 2-mL aliquot of supernatant was transferred into glass vials and concentrated in a vacuum oven (Stegler VAC-52 FCD-3000, Shanghai, China). The dried residues were reconstituted in 1 mL of 40:60 (v/v) acetonitrile/milli-Q water. Extracts underwent ultrasonication for 10 minutes and were transferred to microtubes. 50 µL of 10% trichloroacetic acid solution was added to each sample. Microtubes were vortex-mixed for 1 minute and centrifuged at 16,000 rpm during 10 min (Microspin-12, Biosan Riga, Latvia) (Gonzalez-Rey and Bebianno, 2012). Prior to analysis, samples were filtered through 13-mm PVDF syringe membranes (0.45-µm pore size). A 200-µL aliquot of filtrate was transferred to HPLC vials for analysis.

During the study, three types of samples were measured: amphipod extracts (a), ibuprofen analytical standard solution (Certified Reference Material 11559-2020, NCAS, Moscow, Russia) (b), and amphipod extracts spiked with ibuprofen standard (c). The analytical standard solution was used to optimize ionization parameters and evaluate chromatographic system performance (Eraga et al., 2015). Subsequently, amphipod extracts were analyzed both unmodified and following standard addition. The concentration of the ibuprofen analytical standard working solution was 72 ng/mL. Spiked samples consisted of 600 µL amphipod extract and 100 µL ibuprofen analytical standard solution (final volume: 700 µL) (Bashyal, 2018; Dowling et al., 2020). A linear calibration curve was generated for ibuprofen concentrations ranging from 0,025 ng/mL to 50 ng/mL.

Screening for ibuprofen in Baikal endemic amphipods was performed using an Agilent Infinity II (2019) LC-MS/MS system equipped with an Agilent 6470B triple quadrupole mass spectrometer. Chromatographic separation used an Agilent Poroshell C18 column (2.1 × 50 mm) maintained at 30°C. Table 1 presents the program for HPLC separation of samples containing ibuprofen. Mass spectrometer setup program: ion source gas temperature: 300 °C; ion source gas flow: 5 L/min; nebuliser: 45 psi; drying gas temperature: 250 °C; drying gas flow: 11 L/min; capillary voltage: 3500 V; sample volume: 1 µL. Ibuprofen detection used MRM transition 205.1 → 161.1 (GOST 32881-2014, Moscow, Russia).

**Table 1.** HPLC gradient program for ibuprofen separation.

| **Time, min** | **Solution B**<br>**100% Acetonitrile, %** | **Flow Rate, mL/min** |
| --- | --- | --- |
| 1.0 | 40 | 0.4 |
| 4.0 | 98 | 0.4 |
| 6.0 | 98 | 0.4 |
| 7.0 | 40 | 0.4 |
| 7.5 | 40 | 0.4 |

The study involved analysis of 218 specimens of Baikal amphipods, including *E. verrucosus* (n=180), *O. flavus* (n=20), *E. cyaneus* (n=3), and unidentified amphipods of the genera *Eulimnogammarus* (n=8), *Brandtia* sp. (n=6), and *Pallasea* sp. (n=1). Samples of *E. verrucosus* and *Brandtia* sp. comprised single individuals. Analyses of *O. flavus* and *E. cyaneus* utilized pooled samples containing two individuals each.

The statistical processing was performed in Past software (V4.03) using the Kruskal–Wallis H test (one-way ANOVA on ranks as a non-parametric method) for the analysis of qualitative data. The Bonferroni sequential significance model was used. One-way ANOVA, t-test and F-test were used to analyze quantitative data. The effects of factors “year of sampling” and “wet weight of amphipods” were reported based on two-way ANOVA. To determine which group means were significantly different from each other, Tukey’s post-hoc tests were performed. Differences between the mean values of the parameters were considered significant at p ≤ 0.05.

## RESULTS

### Qualitative assessment of the ibuprofen presence in Baikal endemic amphipod

Qualitative assessment of ibuprofen presence was carried out in amphipods of the species *Eulimnogammarus verrucosus* collected in autumn 2023 and *Ommatogammarus flavus* collected in spring 2023. The qualitative analysis demonstrated that ibuprofen was detected in *E. verrucosus* collected in the Angara River, Listvyanka settl., and Buguldeyka settl. The detection frequency of ibuprofen contamination in amphipods ranged from 12% to 27%. Ibuprofen was not detected in amphipods collected in Bolshoye Goloustnoye, Kultuk, and Ust-Barguzin settlements (Fig. 2).

**Fig. 2.**
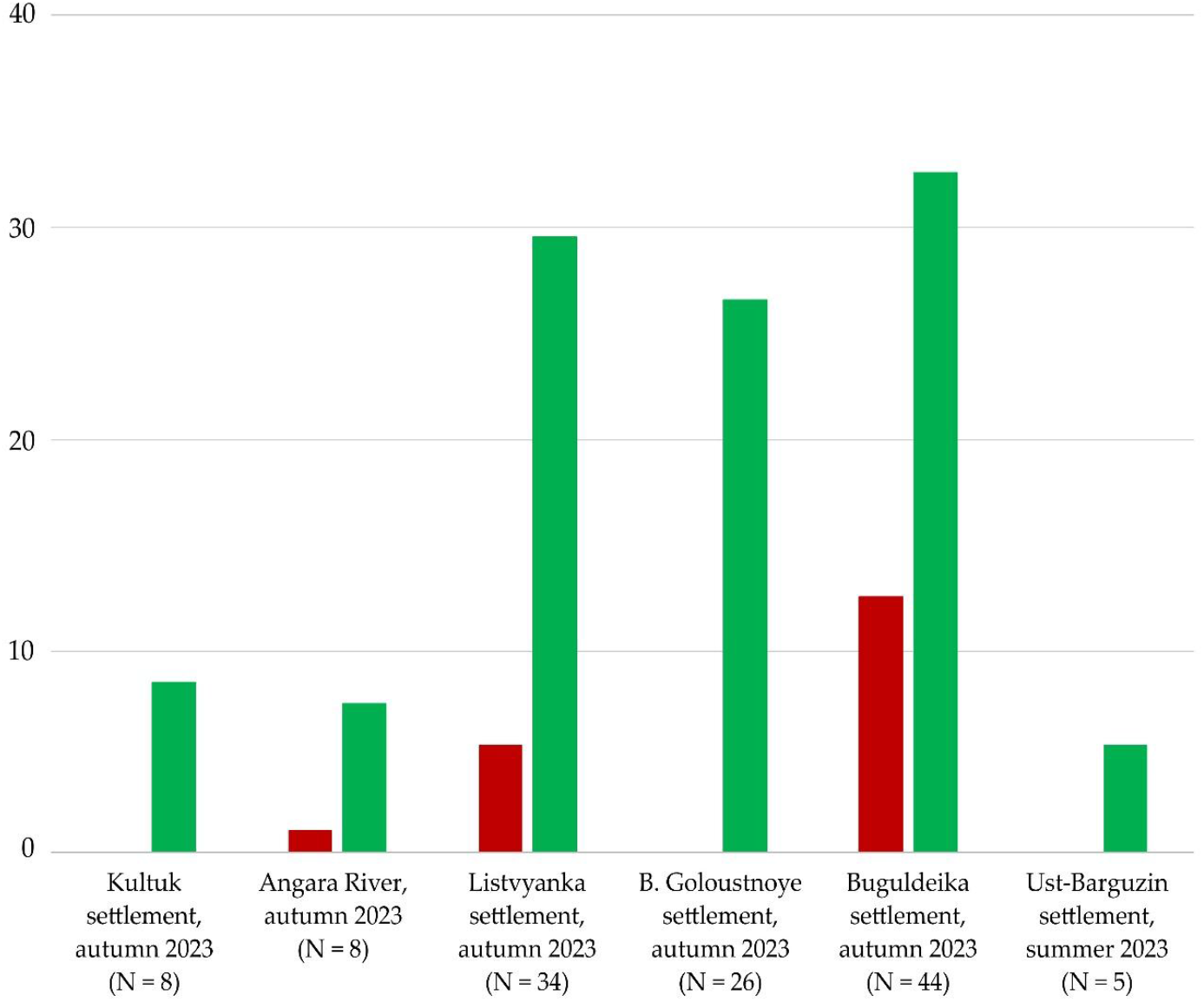
Qualitative assessment of ibuprofen in distinct populations of *E. verrucosus* 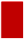 – *E. verrucosus* contaminated with ibuprofen; 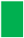 – clean samples

The chromatogram of the ibuprofen analytical standard (Fig. 3a), chromatogram of ibuprofen detected in *E. verrucosus* (Fig. 3b), and chromatogram of amphipod extract spiked with ibuprofen standard (Fig. 3c) demonstrate identical retention times and mass spectra, confirming ibuprofen presence in endemic amphipods.

**Fig. 3.**
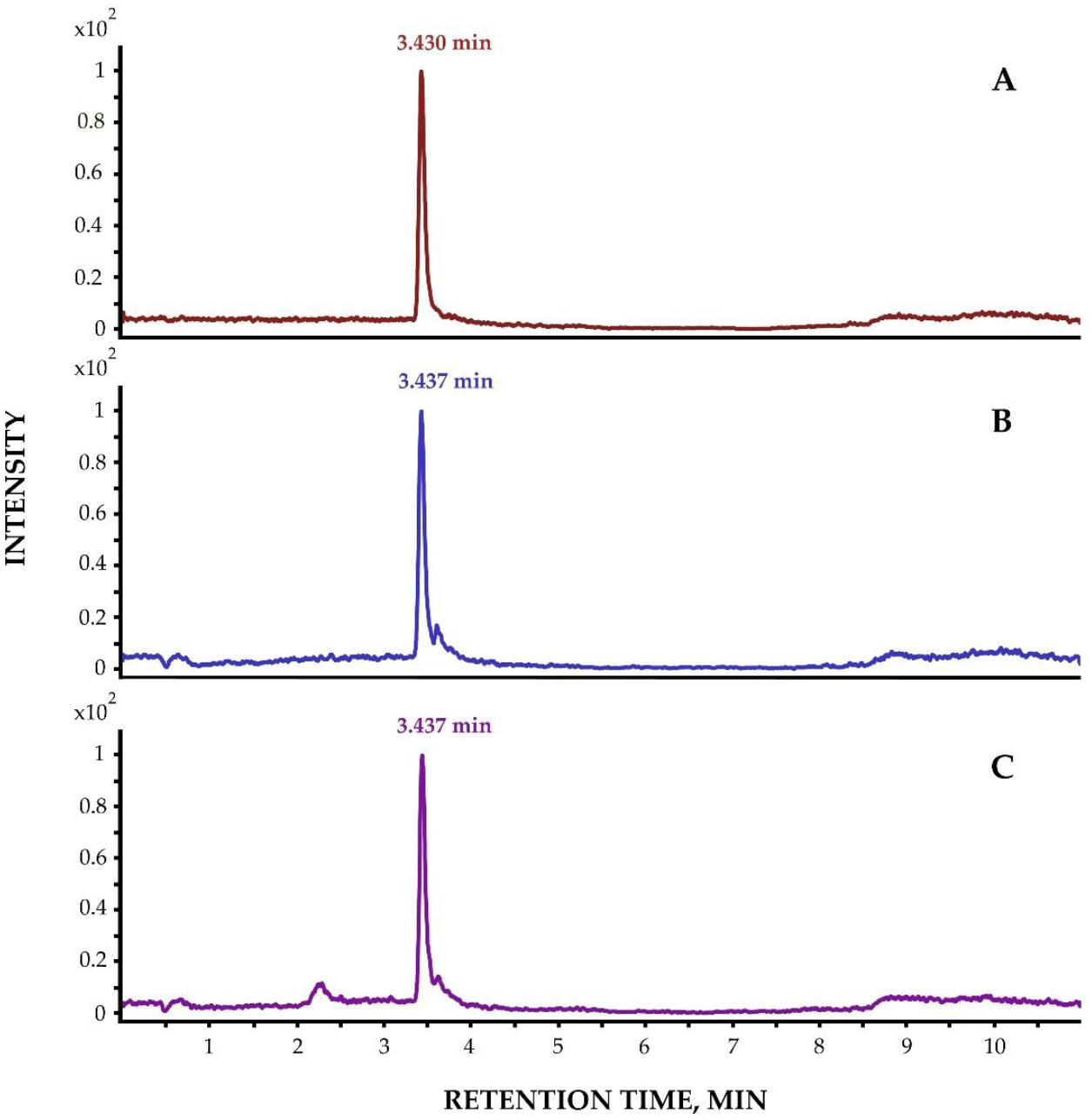
Representative chromatograms: (A) Ibuprofen analytical standard; (B) Ibuprofen detected in *E. verrucosus* extract; (C) *E. verrucosus* extract spiked with ibuprofen standard

Ibuprofen was detected in the deep-water amphipod *O. flavus* collected at 70 m depth near Buguldeika settl. One of 20 analyzed specimens tested positive for ibuprofen. Representative chromatograms are shown in Figure 4. Also, qualitative assessment confirmed ibuprofen presence in *E. cyaneus* and *Pallasea* sp. However, insufficient biomaterial precluded quantification of contamination levels or internal ibuprofen concentrations.

**Fig. 4.**
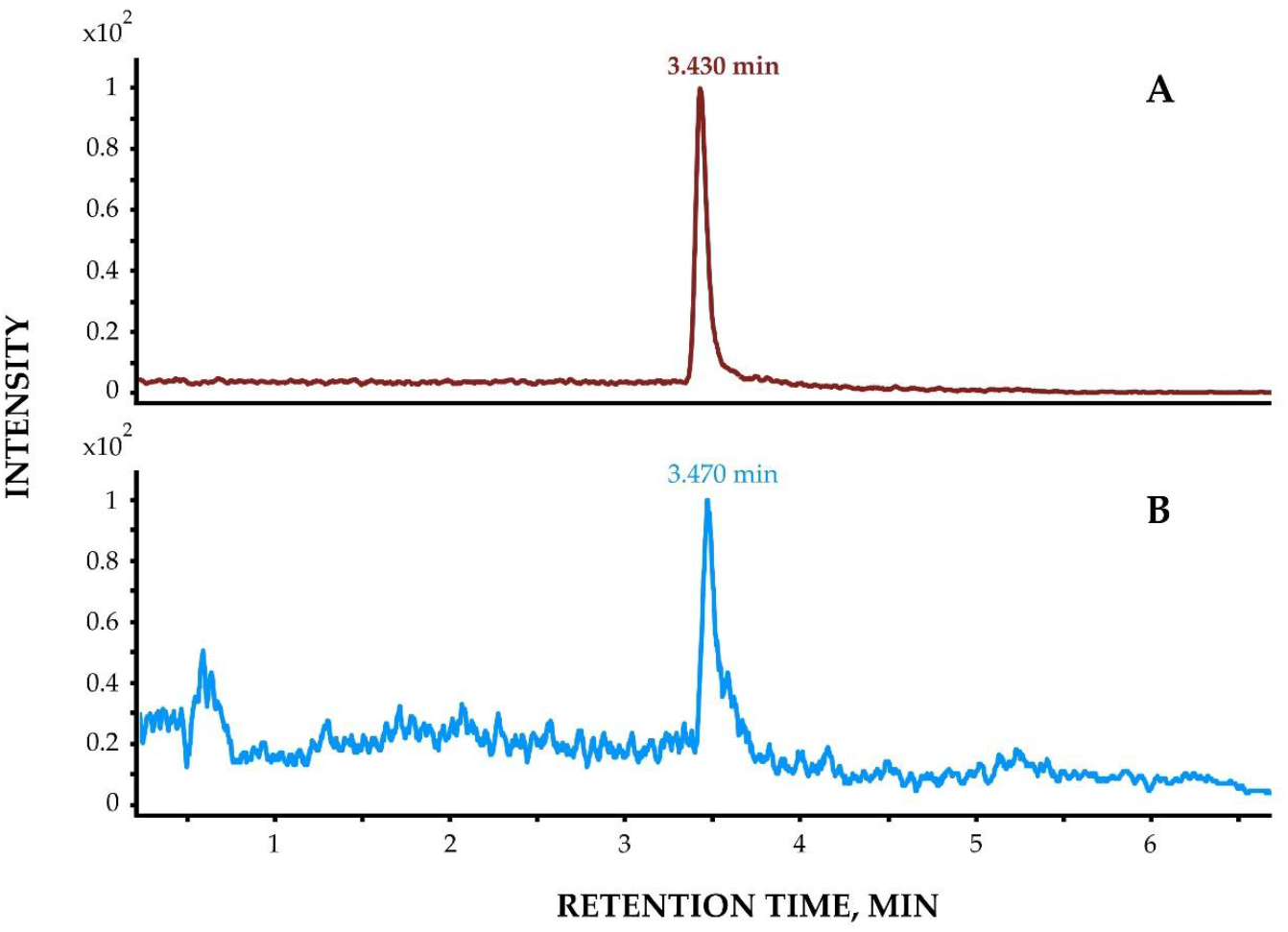
Representative chromatograms of ibuprofen detection: (A) Analytical standard; (B) *O. flavus* extract with detected ibuprofen

### Quantitative assessment of ibuprofen content in Baikal endemic amphipod

Quantitative assessment of ibuprofen in *E. verrucosus* from Listvyanka settl. revealed significantly higher accumulation during spring 2023 compared to autumn 2023 (p = 0.001). Spring amphipods accumulated twice the ibuprofen levels measured in their autumn counterparts (Fig. 5). This seasonal difference indicates dynamic bioaccumulation patterns in Baikal’s endemic amphipods.

**Fig. 5.**
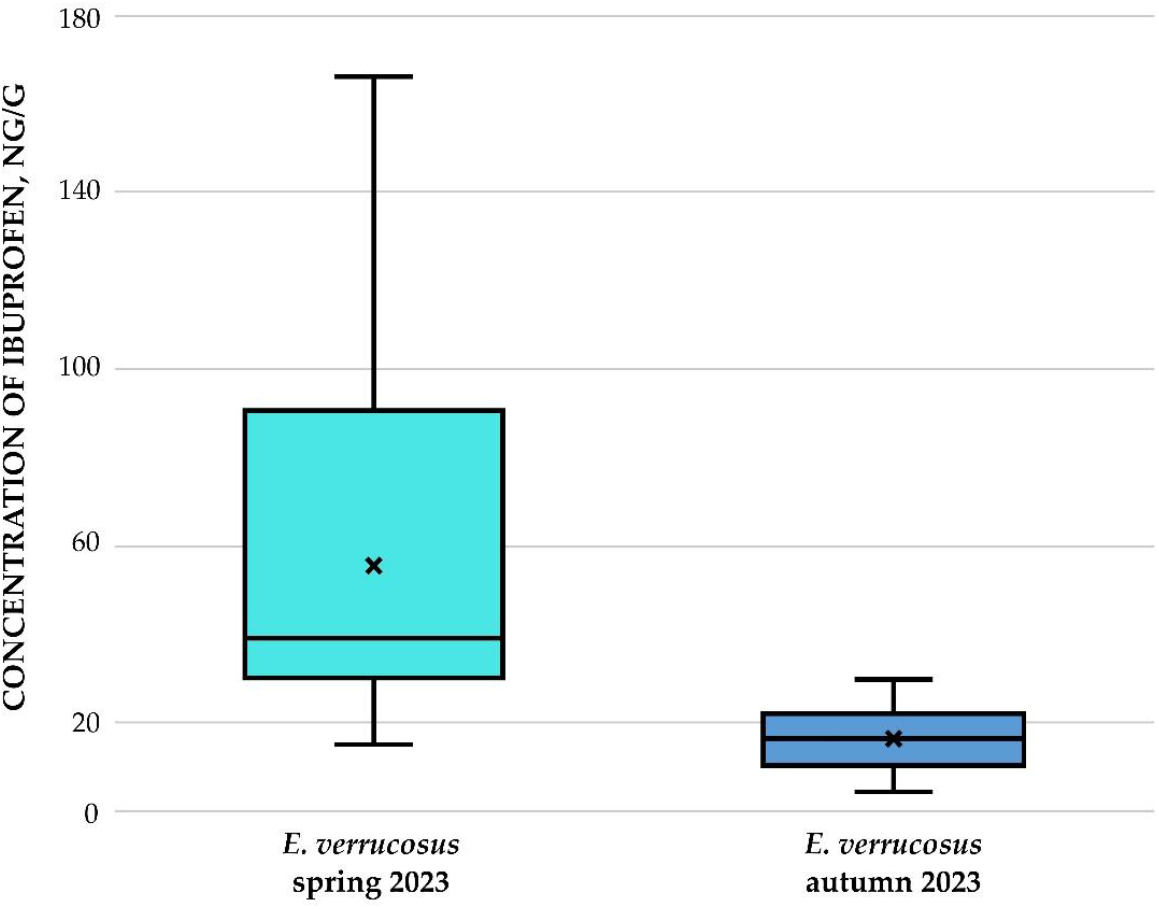
Concentration of ibuprofen (in ng/g) in amphipods of species *E. verrucosus* collected in two seasons in Listvyanka settl. in 2023

In *E. verrucosus* collected during spring, ibuprofen concentrations ranged from 14.92 to 166.38 ng/g, while wet weights ranged from 74 to 776 mg. During autumn, concentrations ranged from 4.19 to 29.65 ng/g, with wet weights spanning 213 to 431 mg.

Amphipods of the genera *Eulimnogammarus* and *Brandtia* collected from the Angara River in spring 2023 were subsequently analyzed. Quantitative assessment revealed significant interspecific differences in ibuprofen accumulation: *Brandtia* specimens accumulated fourfold higher concentrations than *Eulimnogammarus* (p = 0.001; Fig. 6). In amphipods of the genus *Eulimnogammarus*, ibuprofen concentrations ranged from 16.84 to 46.50 ng/g, with individual wet weights spanning 285 mg to 440 mg. For amphipods of the genus *Brandtia*, wet weights ranged from 28 mg to 68 mg, while ibuprofen concentrations ranged from 174.9 to 439.69 ng/g.

**Fig. 6.**
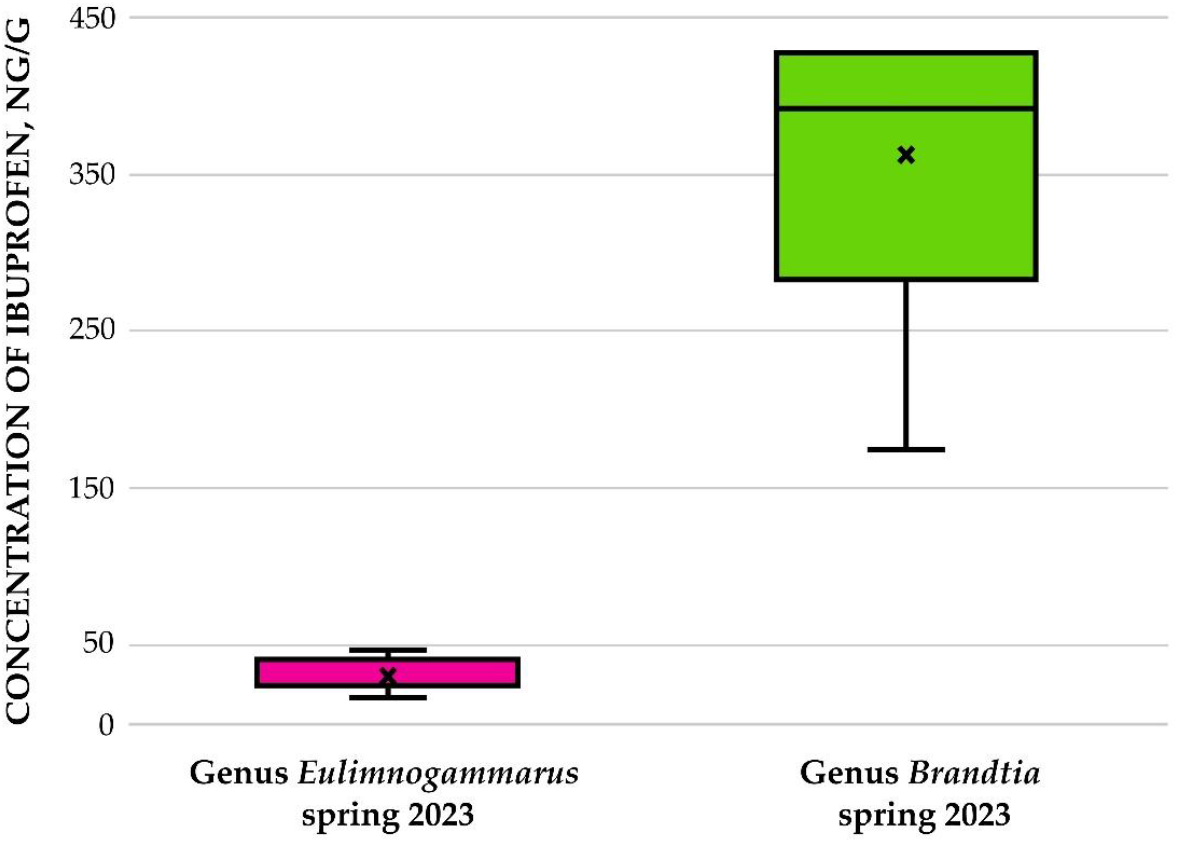
Concentration of ibuprofen (in ng/g) in amphipods of genera *Eulimnogammarus* and *Brandtia*, collected in the Angara River

The maximum ibuprofen concentration was detected in *E. verrucosus* collected during autumn near Buguldeika settl. (Fig. 7). For specimens with wet weights ranging from 105 mg to 307 mg, ibuprofen levels ranged from 18.33 to 1151.32 ng/g. In the studied individual of *Pallasea* sp. (wet weight: 0.130 g), ibuprofen was detected at 84.86 ng/g, while the concentration in deep-water *O. flavus* was 5.15 ng/g.

**Fig. 7.**
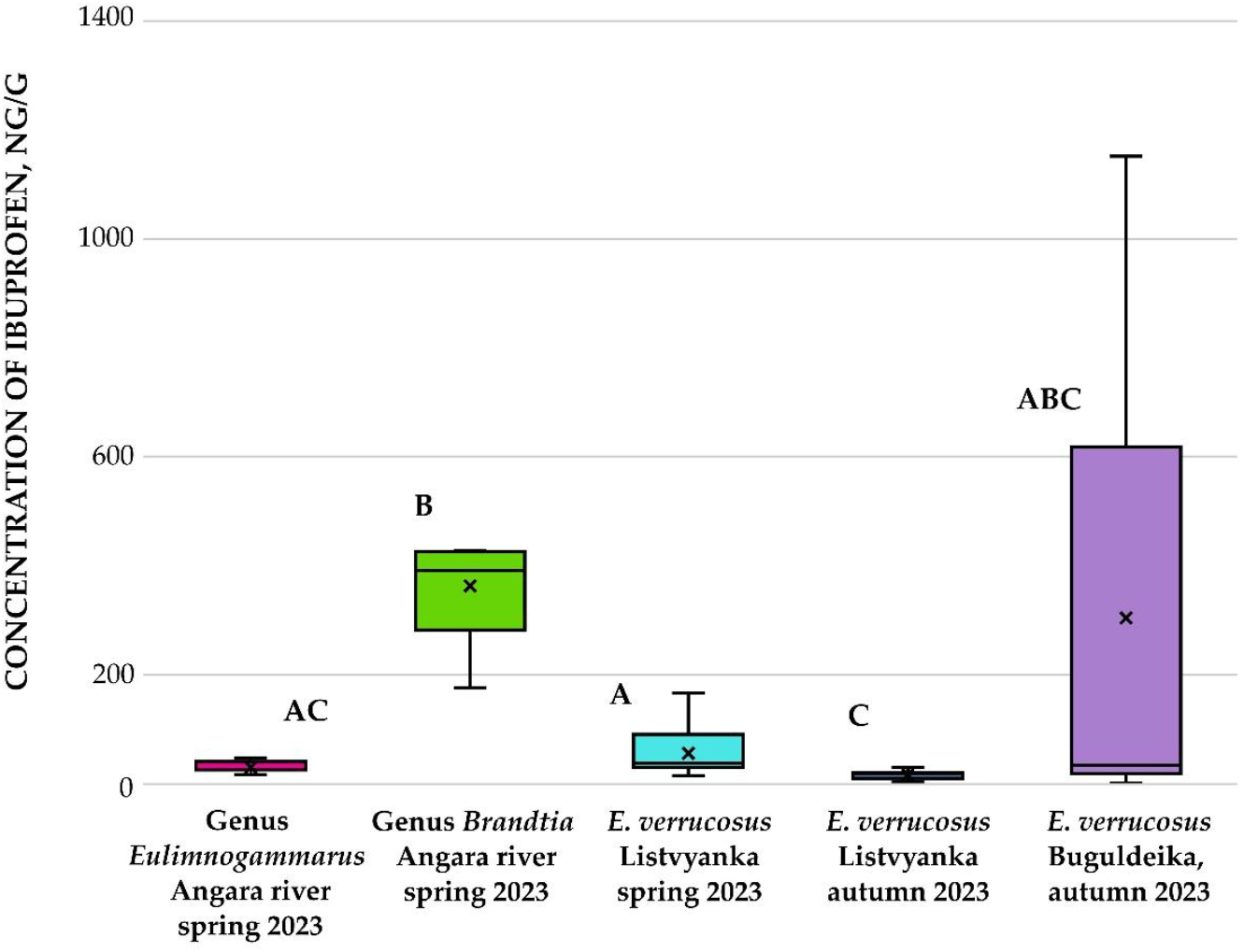
Concentration of ibuprofen (in ng/g) in amphipods collected in two seasons in 2023 Note: letter designations of amphipod groups denote statistically significant differences

Furthermore, a dependence of ibuprofen concentration on organism wet weight was observed in *E. verrucosus* collected during autumn near Listvyanka settl. (Fig. 8). In amphipods with wet weights from 0.21 to 0.29 g, the maximum ibuprofen concentration reached 29.65 ng/g, while specimens weighing 0.43 g contained ibuprofen at 4.19 ng/g. This weight-concentration relationship was also evident in *Brandtia* spp. from the Angara River (Fig. 9), where amphipods weighing 0.028-0.036 g showed concentrations of 385.03-431.63 ng/g, whereas a 0.068 g specimen accumulated ibuprofen at 174.90 ng/g.

**Fig. 8.**
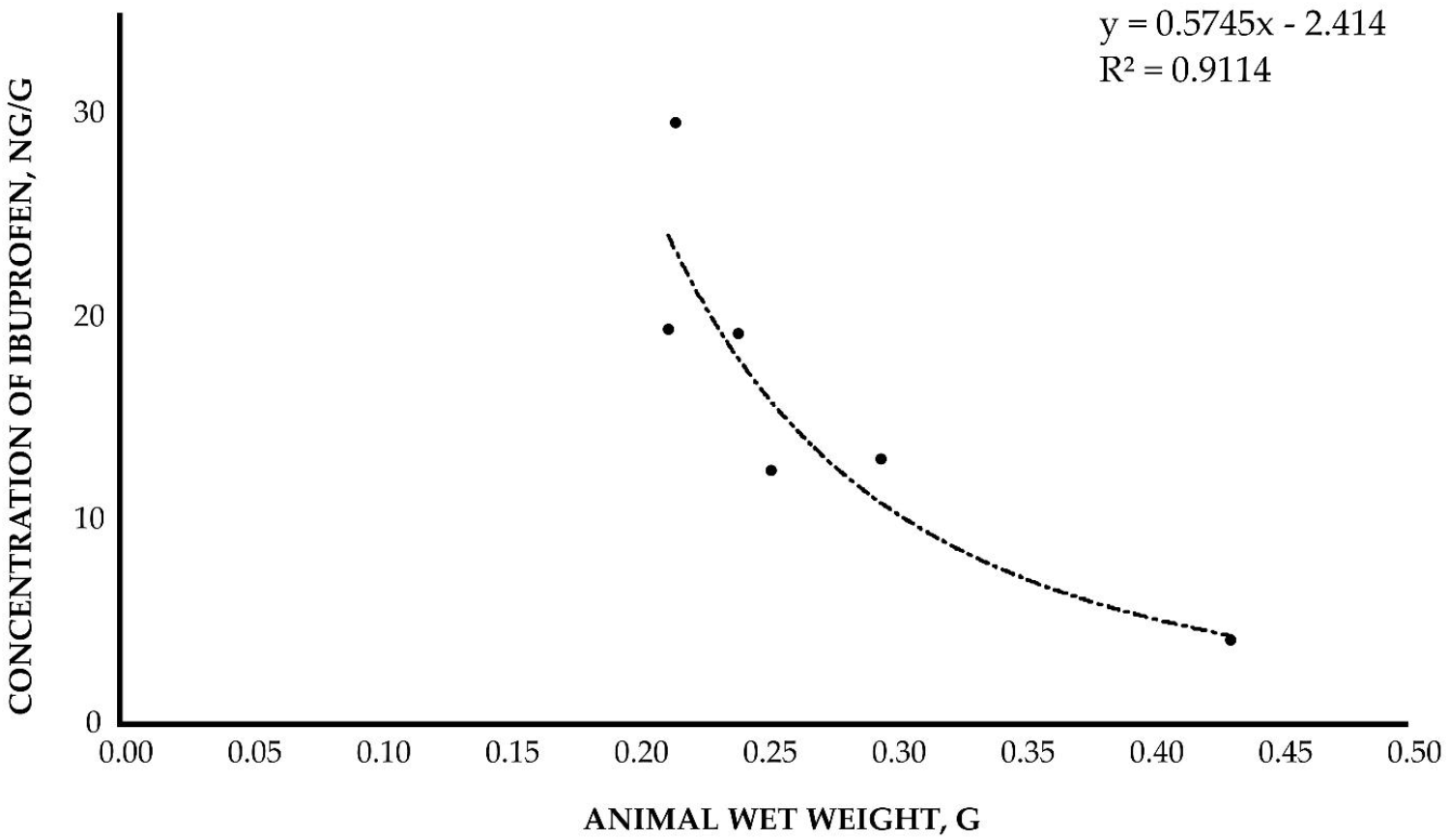
The correlation between the concentration of accumulated ibuprofen and wet weight of amphipod *E. verrucosus* (in ng/g). Sampling site: Listvyanka settl. (autumn 2023)

**Fig. 9.**
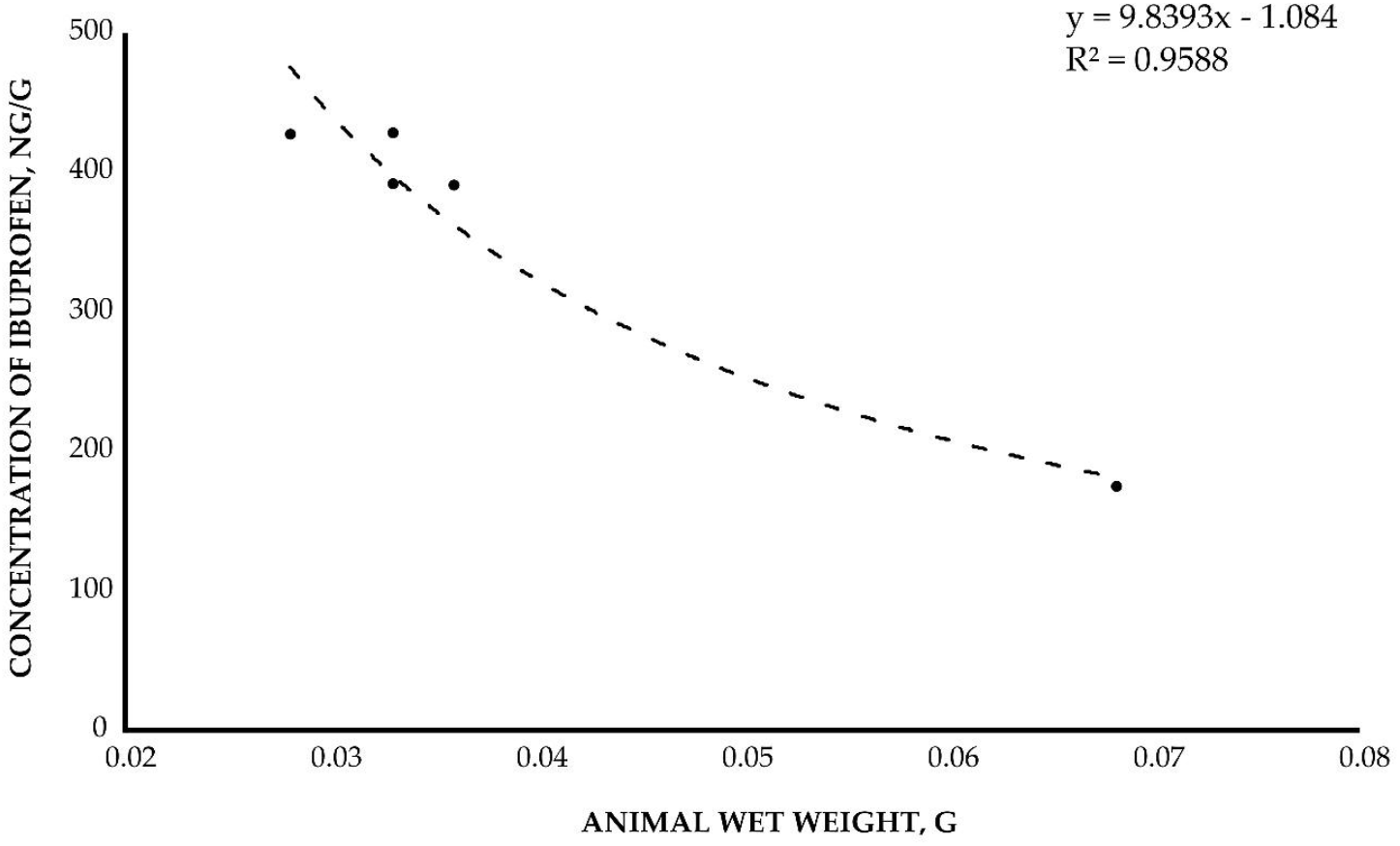
The correlation between the concentration of accumulated ibuprofen and wet weight of amphipod *Brandtia* (in ng/g). Sampling site: Angara River (spring 2023)

Additionally, a compound structurally similar to ibuprofen was detected in several *E. verrucosus* specimens. While ibuprofen shows MRM transition 205.1 → 161.1, this compound exhibited a retention time shift of +0.2 minutes (Fig. 10C) with identical mass transitions. We hypothesize that this substance corresponds to the ibuprofen metabolite, since both substances have a similar fragmentation pattern and a similar retention time. An ibuprofen derivative was detected in *E. verrucosus* collected from the Angara River, Listvyanka, Bolshoye Goloustnoye and Buguldeika settlements (Fig. 11). The detection frequency of this derivative ranged from 10% to 50%. It was not detected in *E. verrucosus* from Kultuk and Ust-Barguzin settlements, nor in the deep-water amphipod *O. flavus*.

**Fig. 10.**
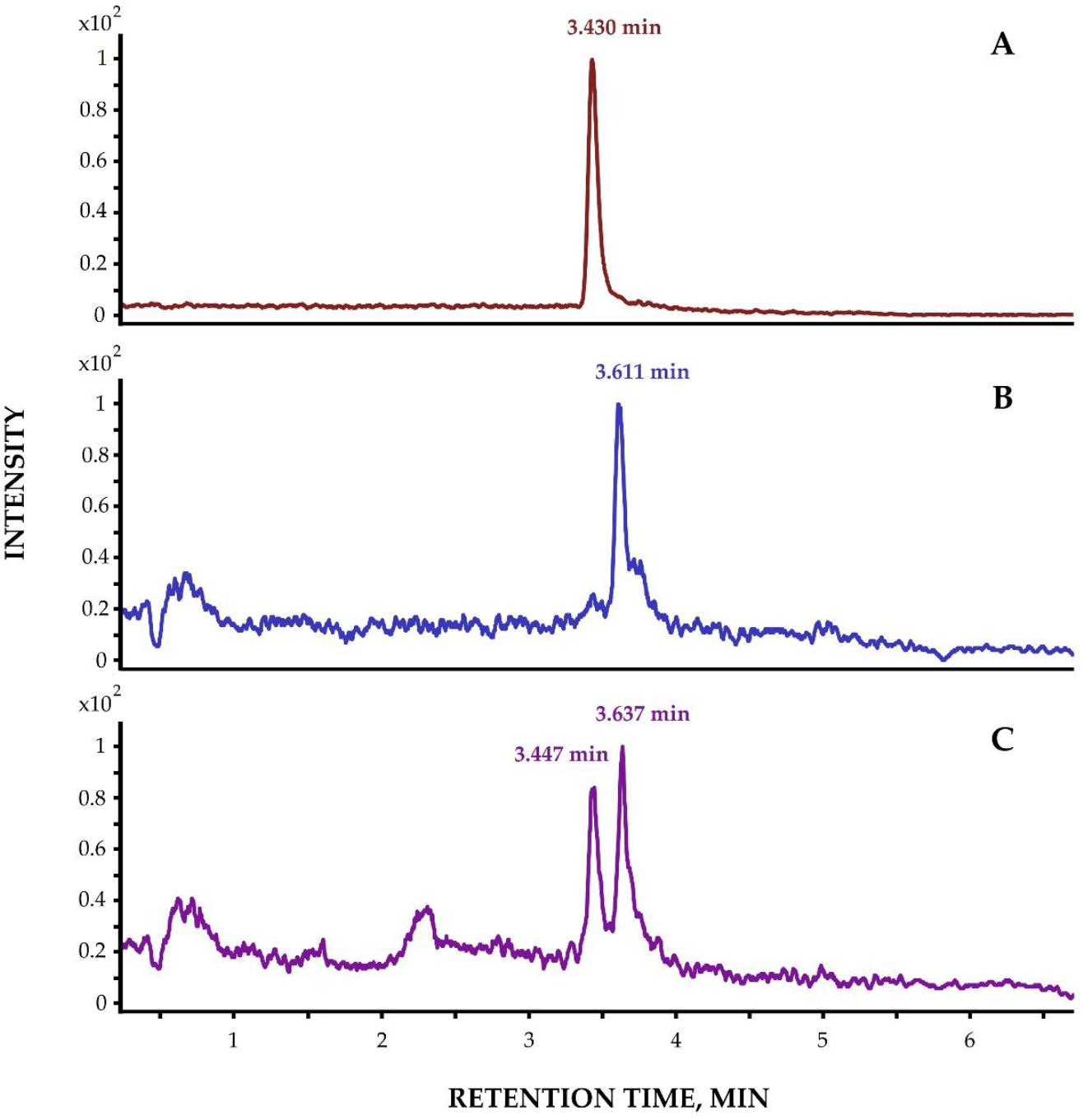
Representative chromatograms: (A) Ibuprofen analytical standard; (B) Suspected ibuprofen metabolite in *E. verrucosus* extract; (C) *E. verrucosus* extract spiked with ibuprofen standard

**Fig. 11.**
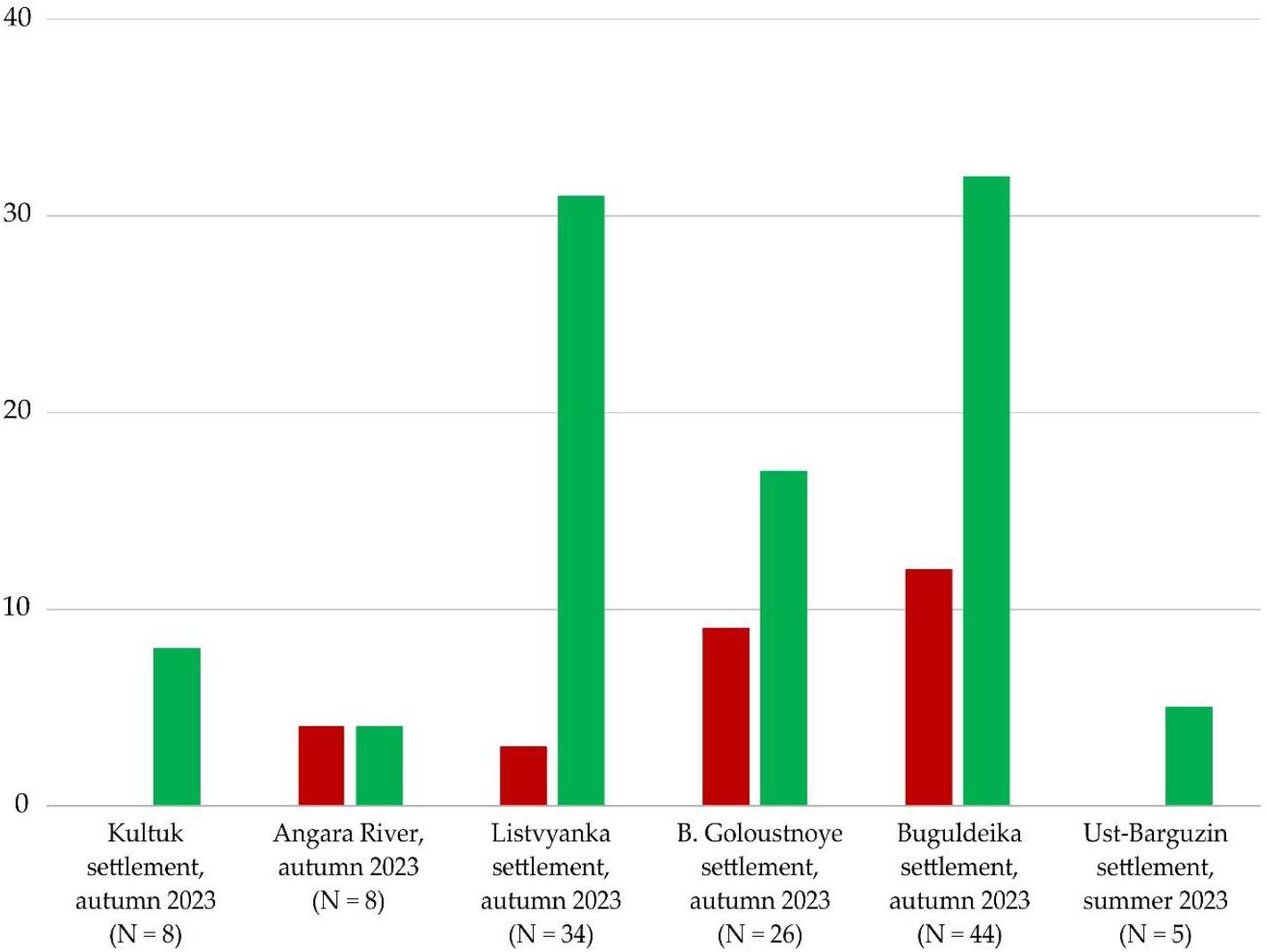
Evaluation of the presence of ibuprofen derivative in different populations of amphipod species *E. verrucosus* 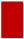 – *E. verrucosus* contaminated with ibuprofen derivative; 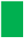 – clean samples

## DISCUSSION

Pharmaceutical bioaccumulation in Lake Baikal’s ecosystem represents an urgent ecological concern. Initial studies confirmed trace drug residues in endemic amphipods, including acetylsalicylic acid, paracetamol, azithromycin, tetracyclines, amikacin, dimetridazole, metronidazole, and spiramycin. Earlier ibuprofen detection was reported by our studies, published by Telnova et al. (2024), who analyzed *E. verrucosus* from a Bolshoe Goloustnoe settl. in August 2020 and 2022. This monitoring revealed significant temporal variation: ibuprofen was detected in 70% of specimens in 2020 versus 27% in 2022. Our 2023 analysis revealed no direct ibuprofen contamination. Here we detected ibuprofen transformation products in 34% of *E. verrucosus* specimens. While ibuprofen metabolite screening was not conducted in 2020-2022, the declining parent compound concentration may reflect reduced post-pandemic pharmaceutical loading. These findings demonstrate that Baikal amphipods experience intermittent, but recurrent, pharmaceutical exposure with shifting contaminant profiles.

Quantitative analysis confirmed a significant inverse relationship between amphipod wet weight and accumulated ibuprofen concentrations. Maximum ibuprofen levels occurred in *E. verrucosus* from Buguldeika settl. Interspecific comparisons of *Eulimnogammarus* sp. and *Brandtia* sp. in the Angara River revealed differential bioaccumulation capacities. These patterns likely reflect morphological adaptations, particularly differences in integument permeability and exoskeleton composition affecting chemical uptake. Supporting this mechanism, Jakob et al. (2017) demonstrated reduced cadmium accumulation in larger *E. verrucosus* versus smaller *E. cyaneus* – attributable to allometric scaling of surface-area-to-volume ratios. Such size- and species-dependent accumulation principles appear conserved across pollutant classes, extending to pharmaceutical contaminants in Baikal amphipods.

Previous studies (Meyer et al., 2022) detected pharmaceuticals in Lake Baikal water including paracetamol (acetaminophen), paraxanthine, caffeine, cotinine, cimetidine, diphenhydramine, phenazone and sulphachloropyridazine at concentrations of 1–64 ng/L (equivalent to 0.001–0.064 ng/g). Contrastingly, our 2023 data reveal substantial bioaccumulation in endemic amphipods, with ibuprofen concentrations ranging from 4.19 to 1151.32 ng/g – representing bioaccumulation factors (BAFs) of 65,500 to 1,151,320 relative to environmental concentrations (Chen et al., 2023). Conservative estimates indicate amphipods concentrate ibuprofen at 4,000–17,000× environmental levels when compared to typical non-zero contaminant measurements (≥1 ng/L). This demonstrates exceptional biomagnification capacity within Baikal’s trophic web.

We propose that littoral *E. verrucosus* metabolizes ibuprofen, evidenced by a consistent 0.2-minute chromatographic retention time shift with identical mass fragmentation (MRM 205.1→161.1), indicating structural analogs. This derivative likely forms via amphipod enzymatic or symbiotic biotransformation. Notably, the metabolite was absent in *E. verrucosus* from Kultuk and Ust-Barguzin settlements.

The accumulation of active pharmaceutical ingredients like ibuprofen in Lake Baikal’s endemic amphipods originates from anthropogenic sources, entering the ecosystem via sewage discharge and groundwater infiltration—either as parent compounds or bioactive metabolites (Timoshkin et al., 2016). As a synthetic xenobiotic, ibuprofen lacks natural emission pathways. This contamination is exacerbated by Baikal’s status as a major recreational hub, where tourism and local usage drive pharmaceutical loading into nearshore environments.

This study identifies Listvyanka settlement as a principal ibuprofen contamination hotspot, where *E. verrucosus* amphipods exhibit dual exposure responses: bioaccumulation of parent compounds and metabolic transformation into derivatives. Anthropogenic loading likely stems from intensive tourism and inadequate wastewater treatment, compounded by seasonal dynamics. Quantitative analysis confirms significantly higher concentrations in spring-collected specimens versus autumn, potentially reflecting elevated pharmaceutical consumption during winter illness peaks.

Hydrographic processes critically govern ibuprofen distribution in Lake Baikal, where the counter-clockwise surface current (0.5–1.8 m/s) transports contaminants southward along the western shore from high-tourism zones (Maloe More and Olkhon Island) toward downstream settlements. This advective flow establishes a contamination gradient: Buguldeika—closest to northern sources (50 km from Olkhon) and directly within the current – exhibited peak concentrations (1151 ng/g); Listvyanka (mid-transit, 85 km downstream) showed moderate levels (166 ng/g); while Kultuk, positioned beyond the Khamar-Daban coastal deflection zone and farthest from sources (120 km), showed no contamination. This spatial pattern correlates with Olkhon Island’s 300% tourism increase since 2015 (Aleksandrova et al., 2021), confirming current-mediated dispersal as the primary distribution mechanism for pharmaceutical pollutants in Baikal’s pelagic ecosystem.

As previously noted, pharmaceutical pollution in Lake Baikal’s waters is significantly driven by the absence of treatment facilities near populated areas and tourist hubs, improper disposal of expired medications, and seasonal disease outbreaks in humans. Critically, no data currently exist on the effects of ibuprofen—or other pharmaceuticals—on amphipods or any endemic Baikal species. Nevertheless, synergistic interactions between pharmaceutical contamination and compounding stressors—including climate change, escalating cyanobacterial proliferation risks, the decline of Baikal sponges, and tourism pressures on sites like Olkhon Island—threaten to accelerate biodiversity loss and facilitate invasive species establishment in the near future (Timoshkin et al., 2016; Moore et al., 2009, 2019). Ongoing monitoring efforts have accumulated a substantial body of data, underscoring both the escalating environmental threat and the critical need for sustained surveillance and proactive conservation measures.

## CONCLUSIONS

This study establishes that endemic littoral and deep-water amphipods in Lake Baikal exhibit sustained ibuprofen contamination. We demonstrate for the first time that amphipods—potentially aided by symbiotic microorganisms—metabolize ibuprofen. Furthermore, Baikal amphipods bioaccumulate this pharmaceutical compound, with significantly higher concentrations observed in spring compared to autumn. Bioaccumulation capacity correlates positively with amphipod body mass, while several populations remain uncontaminated despite proximity to pollution sources. Collectively, these findings confirm that wild populations of endemic Baikal amphipods are chronically exposed to ibuprofen within their natural habitat.

## Ethical Approval

This is not applicable

## Consent to Participate

This is not applicable

## Consent to Publish

This is not applicable

## REFERENCES

Aleksandrova A. Y., Bobylev S. N., Solovyeva S. V., Khovavko I. Y. 2021. Overtourism at Baikal: problems and ways of addressing them. Geography and Natural Resources 3: 248–257. 10.1134/S1875372821030033

Axenov-Gribanov D., Bedulina D., Shatilina Z., Jakob L., Vereshchagina K., Lubyaga Y., Gurkov A., Shchapova E., Luckenbach T., Lucassen M., Sartoris F. J., Pörtner H. O., Timofeyev M. 2016. Thermal preference ranges correlate with stable signals of universal stress markers in Lake Baikal endemic and Holarctic amphipods. PloS one 10: e0164226. 10.1371/journal.pone.0164226

Bashyal S. 2018. Ibuprofen and its different analytical and manufacturing methods: A review. Asian Journal of Pharmaceutical and Clinical Research 11: 25–29. 10.22159/ajpcr.2018.v11i7.24484

Berlioz-Barbier A., Bulete A., Fabure J., Garric J., Cren-Olive C., Vulliet E. 2014. Multi-residue analysis of emerging pollutants in benthic invertebrates by modified micro-quick-easy-cheap-efficient-rugged-safe extraction and nanoliquid chromatography–nanospray–tandem mass spectrometry analysis. Journal of Chromatography A 1367: 16–32. 10.1016/j.chroma.2014.09.044

Borges K. B., de Oliveira A. R. M., Barth T., Jabor V. A. P., Pupo M. T., Bonato P. S. 2011. LC– MS–MS Determination of ibuprofen, 2-hydroxyibuprofen enantiomers, and carboxyibuprofen stereoisomers for application in biotransformation studies employing endophytic fungi. Analytical and Bioanalytical Chemistry 399: 915–925. 10.1007/s00216-010-4329-9

Chen X., Qadeer A., Liu M., Deng L., Zhou P., Mwizerwa I. T., Jiang X. 2023. Bioaccumulation of emerging contaminants in aquatic biota: PFAS as a case study. Emerging Aquatic Contaminants, Elsevier 347–374. 10.1016/B978-0-323-96002-1.00010-9

Das S. A., Karmakar S., Chhaba B., Rout S. K. 2019. Ibuprofen: its toxic effect on aquatic organisms. Journal of Experimental Zoology India 2: 1125–1131. https://www.researchgate.net/publication/364209489

Ding T., Li W., Li J. 2019. Toxicity and metabolic fate of the fungicide carbendazim in the typical freshwater diatom Navicula species. Journal of Agricultural and Food Chemistry 24: 6683–6690. 10.1021/acs.jafc.8b06179

Dowling G., Gallo P., Fabbrocino S., Serpe L., Regan L. 2008. Determination of ibuprofen, ketoprofen, diclofenac and phenylbutazone in bovine milk by gas chromatography-tandem mass spectrometry. Food Additives and Contaminants. Part A 12: 1497–1508. 10.1080/02652030802383160

Du J., Mei C. F., Ying G. G., Xu M. Y. 2016. Toxicity thresholds for diclofenac, acetaminophen and ibuprofen in the water flea Daphnia magna. Bulletin of Environmental Contamination and Toxicology 97: 84–90. 10.1007/s00128-016-1806-7

Drozdova P., Shatilina Z., Mutin A., Saranchina A., Gurkov A., Timofeyev M. 2025. The Curious Case of Eulimnogammarus cyaneus (Dybowsky, 1874): Reproductive Biology of a Widespread Endemic Littoral Amphipod From Lake Baikal. Journal of Experimental Zoology Part A: Ecological and Integrative Physiology 2: 285–293. 10.1002/jez.2891

Eraga S. O., Arhewoh M. I., Chibuogwu R. N., Iwuagwu M. A. 2015. A Comparative UV−HPLC analysis of ten brands of ibuprofen tablets. Asian Pacific Journal of Tropical Biomedicine 10: 880–884. 10.1016/j.apjtb.2015.06.005

Geiger E., Hornek-Gausterer R., Saçan M. T. 2016. Single and mixture toxicity of pharmaceuticals and chlorophenols to freshwater algae Chlorella vulgaris. Ecotoxicology and Environmental Safety 129: 189–198. 10.1016/j.ecoenv.2016.03.032

Gonzalez-Rey M., M. J. Bebianno. 2012. Does non-steroidal anti-inflammatory (NSAID) ibuprofen induce antioxidant stress and endocrine disruption in mussel Mytilus galloprovincialis? Environmental Toxicology and Pharmacology 2: 361–371. 10.1016/j.etap.2011.12.017

GOST 32881-2014; Food products, food raw materials. Method for determination of the non-steroidal anti-inflammatory drug residue content by high performance liquid chromatography—mass spectrometry. Rosstandart: Moscow, Russia, 2016. (In Russian)

Grzesiuk M., Wacker A., Spijkerman E. 2016. Photosynthetic sensitivity of phytoplankton to commonly used pharmaceuticals and its dependence on cellular phosphorus status. Ecotoxicology 25: 697–707. 10.1007/s10646-016-1628-8

Huerta B., Jakimska A., Llorca M., Ruhí A., Margoutidis G., Acuña V., Sabater S., Rodriguez-Mozaz S., Barcelò D. 2015. Development of an extraction and purification method for the determination of multi-class pharmaceuticals and endocrine disruptors in freshwater invertebrates. Talanta 132: 373–381. 10.1016/j.talanta.2014.09.017

Jakob L., Bedulina D. S., Axenov-Gribanov D. V., Ginzburg M., Shatilina Z. M., Lubyaga Y. A., Madyarova E. V., Gurkov A. N., Timofeyev M. A., Pörtner H. O., Sartoris F. J., Altenburger R., Luckenbach T. 2017. Uptake kinetics and subcellular compartmentalization explain lethal but not sublethal effects of cadmium in two closely related amphipod species. Environmental Science & Technology 51: 7208–7218. 10.1021/acs.est.6b06613

Jan-Roblero J., Cruz-Maya J. A. 2023. Ibuprofen: Toxicology and biodegradation of an emerging contaminant. Molecules 28: 2097. 10.3390/molecules28052097

Jurado A., Vázquez-Suñé E., Pujades E. 2021. Urban groundwater contamination by non-steroidal anti-inflammatory drugs. Water 13: 720. 10.3390/w13050720

Kim J. W., Jang H. S., Kim J. G., Ishibashi H., Hirano M., Nasu K., Ichikawa N., Takao Y., Shinohara R., Arizono K. 2009. Occurrence of pharmaceutical and personal care products (PPCPs) in surface water from Mankyung River, South Korea. Journal of Health Science 55: 249–258. 10.1248/jhs.55.249

Larsen C., Yu Z. H., Flick R., Passeport E. 2019. Mechanisms of pharmaceutical and personal care product removal in algae-based wastewater treatment systems. Science of the Total Environment 695: 133772. 10.1016/j.scitotenv.2019.133772

Luo Y., Guo W., Ngo H. H., Nghiem L. D., Hai F. I., Zhang J., Liang S., Wang X. C. 2014. A review on the occurrence of micropollutants in the aquatic environment and their fate and removal during wastewater treatment. Science of the Total Environment 473–474: 619–641. 10.1016/j.scitotenv.2013.12.065

Marchlewicz A., Guzik U., Wojcieszyńska D. 2015. Over-the-counter monocyclic non-steroidal anti-inflammatory drugs in environment—sources, risks, biodegradation. Water, Air, & Soil Pollution 226: 355. 10.1007/s11270-015-2622-0

Matz V. D., Efimova I. M. 2017. Geological history of Baikal. Nature 1: 25–32. (In Russian)

Meyer M. F., Ozersky T., Woo K. H., Shchapov K., Galloway A. W., Schram J. B., Snow D. D., Timofeyev M. A., Karnaukhov D. Y., Brousil M. R., Hampton S. E. 2021. A unified dataset of co-located sewage pollution, periphyton, and benthic macroinvertebrate community and food web structure from Lake Baikal (Siberia). Limnology and Oceanography Letters 7: 62–79. 10.17605/OSF.IO/9TA8Z

Moore M. V., De Stasio Jr. B. T., Huizenga K. N., Silow E. A. 2019. Trophic coupling of the microbial and the classical food web in Lake Baikal, Siberia. Freshwater Biology 64: 138–151. 10.1111/fwb.13201

Moore M. V., Hampton S. E., Izmest’eva L. R., Silow E. A., Peshkova E. V., Pavlov B. K. 2009. Climate change and the world’s «sacred sea» — Lake Baikal, Siberia. BioScience 59: 405–417. 10.1525/bio.2009.59.5.8

Nantaba F., Wasswa J., Kylin H., Palm W. U., Bouwman H., Kümmerer K. 2020. Occurrence, distribution, and ecotoxicological risk assessment of selected pharmaceutical compounds in water from Lake Victoria, Uganda. Chemosphere 239: 124642. 10.1016/j.chemosphere.2019.124642

Nieto E., Corada-Fernández C., Hampel M., Lara-Martín P. A., Sánchez-Argüello P., Blasco J. 2017. Effects of exposure to pharmaceuticals (diclofenac and carbamazepine) spiked sediments in the midge, Chironomus riparius (Diptera, Chironomidae). Science of the Total Environment 609: 715–723. 10.1016/j.scitotenv.2017.07.171

Pajić N. B., Mureškić I., Jevđenić B., Račić A., Gatarić B. 2023. Interchangeability of medications and biopharmaceutical implication of taking drugs with fluids other than water: ibuprofen case study. Brazilian Journal of Pharmaceutical Sciences 59: e22725. 10.1590/s2175-97902023e22725

Parolini M. 2020. Toxicity of the Non-Steroidal Anti-Inflammatory Drugs (NSAIDs) acetylsalicylic acid, paracetamol, diclofenac, ibuprofen and naproxen towards freshwater invertebrates: A review. Science of the Total Environment 740: 140043. 10.1016/j.scitotenv.2020.140043

Puangpetch A., Limrungsikul A., Prommas S., Rukthong P., Sukasem C. 2020. Development and validation of a liquid chromatography-tandem mass spectrometry method for determination of ibuprofen in human plasma. Clinical Mass Spectrometry 15: 6–12. 10.1016/j.clinms.2019.10.002

Rusinek O. T., Takhteev V. V., Gladkochub D. P., Khodzher T. V., Budnev N. M. 2012. Baikal Science. Nauka Publishers: Moscow, Russia. (In Russian)

Takhteev V. 2019. On the current state of taxonomy of the Baikal Lake amphipods (Crustacea: Amphipoda) and the typological ways of constructing their system. Arthropoda Selecta 28: 374–402. 10.15298/arthsel.28.3.09

Telnova T. Y., Morgunova M. M., Shashkina S. S., Vlasova A. A., Dmitrieva M. E., Shelkovnikova V. N., Malygina E. V., Imidoeva N. A., Belyshenko A. Y., Konovalov A. S., Misharina E. A., Axenov-Gribanov D. V. 2024. Detection of pharmaceutical contamination in amphipods of Lake Baikal by the HPLC-MS method. Antibiotics 13: 738. 10.3390/antibiotics13080738

Timoshkin O. A., Samsonov D. P., Yamamuro M., Moore M. V., Belykh O. I., Malnik V. V., Sakirko M. V., Shirokaya A. A., Bondarenko N. A., Domysheva V. M., Fedorova G. A., Kochetkov A. I., Kuzmin A. V., Lukhnev A. G., Medvezhonkova O. V., Nepokrytykh A. V., Pasynkova E. M., Poberezhnaya A. E., Potapskaya N. V., Rozhkova N. A., Sheveleva N. G., Tikhonova I. V., Timoshkina E. M., Tomberg I. V., Volkova E. A., Zaitseva E. P., Zvereva Yu. M., Kupchinsky A. B., Bukshuk N. A. 2016. Rapid ecological change in the coastal zone of lake Baikal (East Siberia): is the site of the world’s greatest freshwater biodiversity in danger? Journal of Great Lakes Research 42: 487–497. 10.1016/j.jglr.2016.02.011

Tyumina E. A., Bazhutin G. A., Cartagena Gómez A. P., Ivshina I. B. 2020. Nonsteroidal antiinflammatory drugs as emerging contaminants. Microbiology 89: 148–163. 10.1134/S0026261720020125

Shirokova Y., Telnes E., Mutin A., Rzhechitskiy Y., Shatilina Z., Sokolova I., Timofeyev M. 2025. Metabolic responses to thermal ramping in two endemic eurybathic amphipods of the genus Ommatogammarus from ancient Lake Baikal. Comparative Biochemistry and Physiology Part A: Molecular & Integrative Physiology 299: 111881. 10.1016/j.cbpa.2025.111881

Vulliet E., Cren-Olivé C., Grenier-Loustalot M. F. 2011. Occurrence of pharmaceuticals and hormones in drinking water treated from surface waters. Environmental Chemistry Letters 9: 103–114. 10.1007/s10311-009-0253-7

Wang C., Wang Y., Xie H., Zhan C., He X., Liu R., Hu R., Shen J., Jia Y. 2022. Establishment and validation of an sil-is lc–ms/ms method for the determination of ibuprofen in human plasma and its pharmacokinetic study. Biomedical Chromatography 36: e5287. 10.1002/bmc.5287

Yang H., Lu G., Yan Z., Liu J., Dong H., Bao X., Zhang X., Sun Y. 2020. Residues, bioaccumulation, and trophic transfer of pharmaceuticals and personal care products in highly urbanized rivers affected by water diversion. Journal of Hazardous Materials 391: 122245. 10.1016/j.jhazmat.2020.122245

Zanuri N. B. M., Bentley M. G., Caldwell G. S. 2017. Assessing the impact of diclofenac, ibuprofen and sildenafil citrate (Viagra®) on the fertilisation biology of broadcast spawning marine invertebrates. Marine Environmental Research 127: 126–136. 10.1016/j.marenvres.2017.04.005

